# Vegetation characteristics control sediment and nutrient retention on but not underneath vegetation in floodplain meadows

**DOI:** 10.1101/2021.05.21.445106

**Authors:** Lena Kretz, Elisabeth Bondar-Kunze, Thomas Hein, Ronny Richter, Christiane Schulz-Zunkel, Carolin Seele-Dilbat, Fons van der Plas, Michael Vieweg, Christian Wirth

**Affiliations:** Systematic Botany and Functional Biodiversity, Life science, Leipzig University, Germany; Helmholtz Centre for Environmental Research (UFZ), Department Conservation Biology, Germany; University of Natural Resources and Life Sciences, Vienna, Institute of Hydrobiology and Aquatic Ecosystem Management, Austria; WasserCluster Lunz, Austria; German Centre for Integrative Biodiversity Research (iDiv) Halle-Jena-Leipzig, Germany; Geoinformatics and Remote Sensing, Institute for Geography, Leipzig University, Germany; Plant Ecology and Nature Conservation, Wageningen University, Droevendaalsesteeg 4, 6708 PB Wageningen, The Netherlands; Max Planck Institute of Biogeochemistry, Germany

## Abstract

Sediment and nutrient retention are essential ecosystem functions that floodplains provide and that improve river water quality. During floods, the floodplain vegetation retains sediment, which settles on plant surfaces and the soil underneath plants. Both sedimentation processes require that flow velocity is reduced, which may be caused by the topographic features and the vegetation structure of the floodplain. However, the relative importance of these two drivers and their key components have rarely been both quantified. In addition to topographic factors, we expect vegetation height and density, mean leaf size and pubescence, as well as species diversity of the floodplain vegetation to increase the floodplain’s capacity for sedimentation. To test this, we measured sediment and nutrients (carbon, nitrogen and phosphorus) both on the vegetation itself and on sediment traps underneath the vegetation after a flood at 24 sites along the River Mulde (Germany). Additionally, we measured biotic and topographic predictor variables. Sedimentation on the vegetation surface was positively driven by plant biomass and the height variation of the vegetation, and decreased with the hydrological distance (total R^2^=0.56). Sedimentation underneath the vegetation was not driven by any vegetation characteristics but decreased with hydrological distance (total R^2^=0.42). Carbon, nitrogen and phosphorus content in the sediment on the traps increased with the total amount of sediment (total R^2^=0.64, 0.62 and 0.84, respectively), while C, N and P on the vegetation additionally increased with hydrological distance (total R^2^=0.80, 0.79 and 0.92, respectively). This offers the potential to promote sediment and especially nutrient retention via vegetation management, such as adapted mowing. The pronounced signal of the hydrological distance to the river emphasises the importance of a laterally connected floodplain with abandoned meanders and morphological depressions. Our study improves our understanding of the locations where floodplain management has its most significant impact on sediment and nutrient retention to increase water purification processes.

## Introduction

Worldwide, streams and rivers suffer from large loads of sediment and nutrients, which is predominantly caused by anthropogenic activities (1–3). Soil erosion and overfertilization, caused by industrial agriculture and forestry, increase the loads of sediment and nutrients in river systems and cause eutrophication and siltation (4–6). Additionally, the process of sediment transport along the river is often interrupted by hydro-engineering infrastructure (6). River floodplains, however, can act as a sink for sediment and its associated nutrients by retaining these during floods (7,8), thus providing the important ecosystem function of sediment and nutrient retention (9,10).

Natural floodplains reduce sediment and nutrient transport to downstream areas during inundation. Especially in hydrologically connected systems, a large amount of the annual riverine sediment and nutrient load can be retained in floodplains. The amount increases with the inundation duration and the area of inundation (11). The accumulated nutrients can have a positive effect on the productivity of the floodplain vegetation (12). However, anthropogenic activities have strongly diminished floodplain areas, due to channelization, embankments, bank stabilization, and river straightening (7,13,14). Consequently, worldwide floodplains are considered threatened ecosystems (13,14). As a result, floodplain restoration efforts have increased during the last decades. Many countries started programs emphasizing the river-floodplain reconnection for restoring ecological conditions, but also for flood protection. Furthermore, reconnection measures are expected to affect the retention capacity of floodplains (15), but its drivers still need to be better integrated into river and floodplain restoration and management (16). However, to manage floodplains for optimal sediment and nutrient retention, we need to understand how vegetation structure, as well as the composition and diversity of plant communities, affect sedimentation and how these biotic drivers interact with the hydromorphological control.

Sediment retention is a complex phenomenon that depends on different biogeomorphic processes in the floodplain (17). While deposition of coarse sediment is mostly influenced by the topography of the floodplain, the vegetation type and structure influencing fluvial processes and sediment transport (18,19) are most relevant for sedimentation of finer grain sizes (17,20,21). Communities of herbaceous vegetation were more efficient in accumulating fine sediment compared to shrublands and floodplain forests (22), and reed beds caused more nitrogen and phosphorus deposition than grass and woodlands (12). Within a flume experiment, we showed in a previous study, that the structural characteristics of the community (biomass, density, height, structural diversity, and leaf pubescence) increase sedimentation under controlled conditions (23). However, this is the first study that investigates *in situ* measurements of a real flood event by (1) focusing on sedimentation within the vegetation, separating the process of sedimentation on vegetation from the process of sedimentation underneath the vegetation, (2) investigated the role of species diversity, leaf surface structure and community structure, and (3) combined these vegetation characteristics with topographical parameters of the floodplain, thus allowing to quantify the relative importance of vegetation and topography.

The sediment retention capacity of a floodplain is known to vary with different structural parameters of the vegetation, mostly measured around (in front and behind) vegetation patches. Generally, it was found that biomass increases sediment retention (20,24,25), which was also the case in the flume experiments for sedimentation on the vegetation (23,26) and partly also underneath the vegetation (23). Dense floodplain vegetation has been suggested to be very efficient in accumulating fine sediment (22,27). It reduces the flow velocity and thus allows sediment to sink and deposit (28,29). Here, also the variation of the vegetation height may have an impact on sedimentation, since varying vegetation height cause turbulence and might increase and decrease flow velocities locally. In the flume experiment a negative relationship was found between height variation and sedimentation on the vegetation (23). It was found that the deposition of finer sediment (silt and clay) is controlled by vegetation height in herbaceous floodplain vegetation (30).

Riparian zones and floodplain meadows are hotspots of biodiversity (14). At the same time, they are one of the most threatened habitats in the world (31,32). Despite this, species diversity *per se* is rarely studied in the context of sediment retention on floodplains, even though it is known to determine other ecosystem functions such as productivity and nutrient dynamics (33). The results of the flume experiment only showed evidence for effects of species richness on sedimentation in the absence of identity effects (26). Species diversity has also been shown to correlate with structural diversity of vegetation (34), which was found to increase sedimentation (23). Dedicated biodiversity experiments have revealed that diverse grasslands exploit the growing space in a complementary fashion and thus have a higher density and taller stature than less diverse grasslands (35,36). While we account for these two variables directly, there may be additional effects that go beyond the mean characteristics of the vegetation. Combining for example tall/sparse with small/dense plant species may be particularly effective for sediment retention. The trait combination might increase the overall sedimentation irrespective of total density or stature. No significant effects of the species diversity of herbaceous vegetation on sediment retention were found in front of, and behind a vegetation patch when comparing monocultures with a three-species mixture in an experiment (37). However, the investigation of a longer diversity gradient under field conditions could yield another picture.

Besides vegetation structure, leaf surface structure of the vegetation also matters for sedimentation. In particular, leaf pubescence has been shown to positively and leaf area on not-haired leaves negatively drive sediment retention at the level of herbaceous leaf surfaces (23,38,39). Therefore, the mean expression of these traits in the vegetation may also be important for sedimentation at the level of floodplain vegetation patches, which has rarely been considered in studies on sedimentation in herbaceous vegetation.

Topographic variables are the main abiotic factors that could explain sediment distribution within the floodplain. Discharge and with it, inundation depth are strongly affected by elevation. It was found that the location within the floodplain is relevant for sedimentation (19). Fine sediment is transported farther along the river and into the floodplain than coarse sediment and only settles in areas with reduced flow velocity (28,29). In general, sedimentation was found to decrease with increasing distance from the river (27,40). However, a straight line does not necessarily represent the topographic diversity of a dynamic riverine floodplain and the winding path the water travels into the floodplain during floods. Therefore, the length of the shortest path of lowest elevation is a better measure of the ways the river water travels from the river into the floodplain during floods. Such a measure for the true ‘hydrological distance’ may thus better represent the topography of the floodplain. Some studies used other terms to describe a similar measure such as the flow path (41,42) or the hydrological connectivity (15,43,44).

The aim of this study was a holistic analysis putting vegetation and topography control in perspective by first disentangling sedimentation on and underneath the vegetation under *in situ* conditions, second by quantifying the relative importance of vegetation characteristics in relation to topographical parameter and third, by investigated the effects of additional vegetation characteristics (species diversity and leaf surface structure) on sedimentation within a vegetation patch. We tested the following hypotheses:

(H1) Sedimentation on and underneath the vegetation increases with increasing vegetation biomass, cover, vertical density, vegetation height and height variation.
(H2) Sedimentation on and underneath the vegetation decreases with increasing hydrological distance from the stream.
(H3) Sedimentation on and underneath the vegetation increases with increasing plant species diversity.
(H4) Sedimentation on the vegetation increases with increasing leaf pubescence and decreasing mean leaf area.
(H5) Total carbon (C), nitrogen (N) and phosphorus (P) in the sediment on and underneath the vegetation increase with the total amount of sediment deposited.

## Material and Methods

### Study site

The study was located along the Mulde River in Central Germany (S1 Figure), close to its mouth into the Elbe River. Along this river section, the river still flows in its natural bed and has been only moderately modified by hydro-engineering infrastructures and bank stabilization in the past. About half of the cut-banks are not embanked. The study took place in the frame of the restoration project ‘Wilde Mulde – Revitalisation of a dynamic riverine landscape in Central Germany ‘. The project area extends between the towns Raguhn and Dessau (51°43’-46’ N, 12°17’-18’ E). Within the project area, we defined three floodplains as study areas in 2016 (S1 Figure). The Mulde River is dammed around 22 km upstream of the project area and has another smaller weir about 5 km upstream of the first study area. Upstream of the study areas, the Mulde River has a mean discharge of 67 m^3^ s^-1^ (gauging station ‘Priorau 560090’). In February 2017 a small flood occurred for several days with overbank flow conditions and with a peak discharge of 353 m^3^ s^-1^ equals a flood with a discharge occurring on average every second year. In general, the study area is a mosaic of hardwood and softwood floodplain forests and meadows, with our study focusing on the floodplain meadows. The topography of the floodplain meadows is strongly formed by the river, creating a mosaic of steep slip-off slopes with gravel banks in front, depressions, and abandoned meanders further away from the river that get reconnected during floods. The dominant species in the meadows are, depending on microtopography and management, *Arrhenatherum elatius, Bromus inermis, Calamagrostis epigejos, Elymus repens* and *Phalaris arundinacea*.

### Vegetation data

In summer 2016, we established a grid of vegetation plots. Within the three study areas, plots were selected to span the elevation gradient of the slip-off slope and the floodplain meadow above mean flow conditions using a stratified random sampling strategy. In autumn 2016 we selected 54 plots (18 plots per study area) for this study using with the following criteria: (i) plots are fully covered by vegetation; (ii) plots span a gradient of vegetation height (ranging from 36 cm to 124 cm); (iii) lower elevation plots were given preference, due to their higher probability to get flooded; (iv) depressions and abandoned meanders at distance to the river were also represented, while ensuring that the selection still represents the whole elevation gradient. With this approach, the plots are representative for the floodplain and at the same time form an observational design by spanning gradients for regression analysis. Within each plot (2 m × 2 m) we identified all vascular plant species and estimated the cover of each species in summer 2016 before the flood. We calculated the Shannon diversity index (45) based on cover. Overall, we inventoried 44 species with the species richness ranging from 2 to 10 species per plot.

### Vegetation characteristics

We measured the maximum height of the vegetation using two metrics: (i) the maximum inflorescence height (highest inflorescence), which represents the maximum vegetation height, and (ii) the maximum canopy height (highest leaf), which represents the maximum height of the vegetation surface. Both metrics were measured with the help of a meter stick five times per plot (in the middle of the square plot and at arm length inside the plot from each corner). We measured the vegetation height at that time point no matter if the vegetation hung over or not. We did this once in summer 2016 before the flood and once in spring 2017 after the flood. Additionally, we took images of side views in the form of cross sections of the vegetation in spring 2017 on all flooded plots to estimate the density and height distribution of the vegetation. To this end, we placed a camera, 1 m with 90° angle in front of the plot (Fig 1). At 50 cm inside the plot we positioned a camera background wall so that every image shows exactly the first 50 cm of the plot (Fig 1, S2 Figure). We carefully pushed down the vegetation outside the plot with a flooring material. Afterwards we analysed the images with the statistical software R (46) for height and density distribution in the same way as done in the flume experiment (23). From these structural images, we derived the variables vertical density, mean height, median height, and height variation (Table 1, S2 Figure). The images were colour normalised and resampled from a resolution of 4000 by 6000 pixels to a resolution of 400 by 600 pixels and afterwards transformed into grey-scale images. In order to perform a binary classification of the image into vegetation and background, we used the otsu-tresholding method (47), as implemented in the package EBImage (48). All variables are described in Table 1.

**Fig 1.**
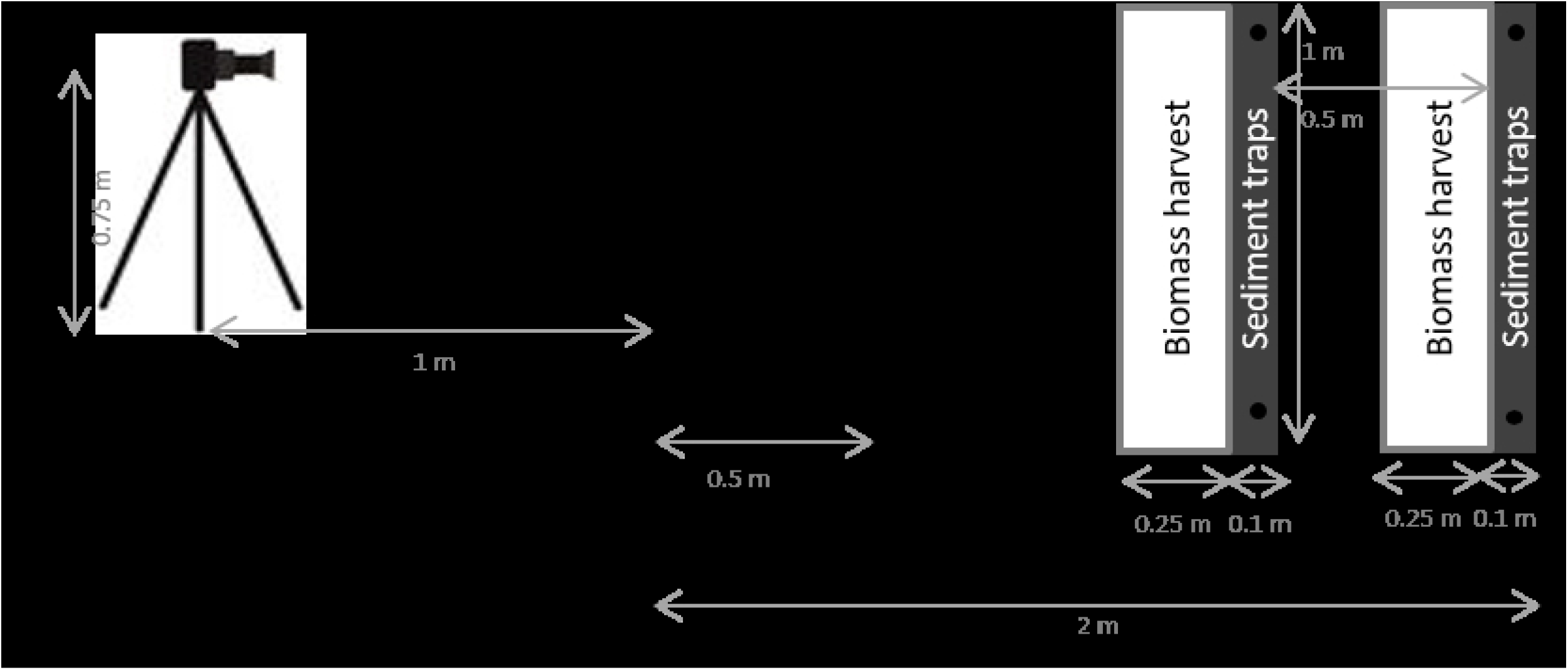
Vegetation plots (2m × 2m). Set-up of the sediment traps and the biomass harvest after the flood event. Set-up of the camera and the camera background for the structural images.

**Table 1:**
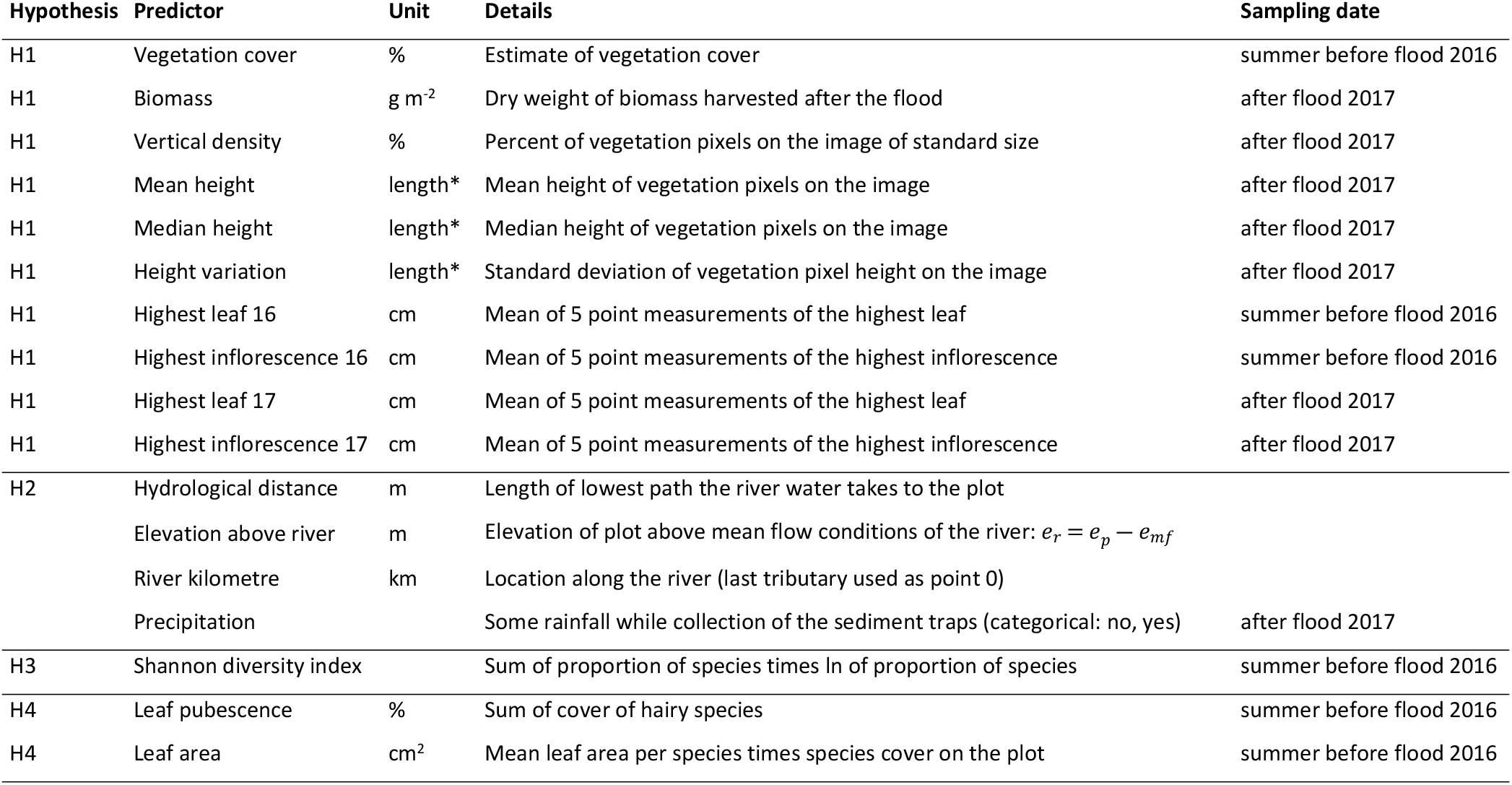
List of predictor variables. Predictor variables with detailed explanations, units and sampling dates. * the length is standardized between the images, however not calibrated to any unit.

### Study design

For investigating sedimentation on the floodplain, we used artificial lawn *(Kunstrasen Arizona*, Hornbach, 1.05 g m^-2^ lawn, 26 cm lawn height, S3 Figure) as sediment traps – a commonly used and established method (29,49). The material has several advantages: (i) it can be easily cut to the required size; (ii) it can be flexibly and firmly fixed to the ground, and (iii) it exposes a surface with a high capacity to collect and keep sediment. To keep the sward structure as intact as possible, we cut the artificial lawn into narrow strips (10 cm × 100 cm strips), which were carefully inserted into the vegetation at two positions within the plot (Fig 1, S3 Figure). While sediment traps represent a good method to measure sedimentation on a standardized surface (thus only affected by surrounding vegetation and its effects on fluvial processes), a limitation is that it removes the effects of the local fine-scale vegetation structure and composition on sedimentation. Combining measures of sedimentation on the vegetation itself, as well as on sediment traps, may be best to partition the effects of fluvial processes (caused by surrounding conditions) and local vegetation properties on sedimentation. We deployed the sediment traps on all 54 plots in January 2017 and fixed them with tent stakes and steel washers (56 cm outer diameter). During the flood in February 2017, 24 plots were inundated (S1 Figure). We collected the sediment traps immediately after the flood retreated. In addition, we also harvested the patch of biomass directly in front of the trap (Fig 1). In the lab, we washed the sediment off the traps with a few litres of water and dried the sediment-rich water in beakers in a compartment drier at 70 °C. Afterwards, the dry sediment was weighed. The same was done with the sediment on vegetation and, additionally, we dried and weighed the biomass itself. The two sediment trap samples per plot were pooled together as were the two biomass samples per plot.

### Nutrient analysis

All sediment samples on the vegetation and on the traps (except two samples with too little sediment) were sieved (< 2 mm) and analysed for C, N and P. To determine the total C and N concentration, the dried sediment samples were ground to a fine powder in a ball mill (Retsch MM2, Vienna, Austria). The homogenized sample was weighed, placed in tin caps and measured by using the Elemental Analysis Isotope Ratio Mass Spectrometry (EA-IRMS; EA—Thermo Scientific™ FLASH 2000 HT™; IRMS-Thermo Scientific™ Delta V™ Advantage) (50). To determine the total P concentration the sediment was also ground to a fine powder in a ball mill (Retsch MM400). The homogenized samples were measured by using the Inductively Coupled Plasma Optical Emission Spectrometry (ICP-OES “Arcos”, Spectro, Kleve, D). As indicator for the nutrient quality the N:P ratio was calculated.

### Topographical variables

The elevation and position of the single plots were measured with a Differential GPS (R8, Trimble Inc.) The mean elevation of the four plot corners *e_p_* was expressed as elevation above the river *e_r_* (as *e_r_* = *e_p_* – *e_mf_*). Mean water level height *e_mf_* was calculated per study area with the digital elevation model (DEM, © GeoBasis-DE, LVermGeo LSA, [m.E. 2016, C22-7009893-2016]) and the closest gauging station (Priorau, 560090). We calculated the elevation difference between the water level of the gauging station on the day the DEM was recorded and the mean water level height (calculated from daily measurements, 1995-2015). With this, we calculated *e_mf_* for each study area. The hydrological distance was defined as the length of the shortest path of lowest elevation that the river water takes to a single plot in the floodplain. It was derived using the flow accumulation approach on the DEM of the floodplain area and calculated using the TopoToolbox 2 (51) in MATLAB (52). We included longitudinal stream distances as river kilometre in the study to account for the plot location along the stream, since we visually observed lower flow velocity at the study area further downstream. The river kilometre was measured along the middle line of the river starting from the last tributary to the river upstream of the study area. We chose this tributary as the zero point because it is the last major tributary. Precipitation occurrence was included as a categorical variable, because some of the traps experienced rainfall after the flood, before all traps could be collected.

### Leaf surface traits

We also included two leaf surface traits, leaf pubescence and leaf area (at plot-level – see below), as predictors of sedimentation, because we showed, with an earlier flume experiment, that, in controlled settings, pubescence can increase leaf surface sedimentation and that sedimentation increases with decreasing leaf area on leaves with no or just a few hairs (39). Out of the 44 species, we classified five as pubescent species (*Carex hirta, Galium aparine, Urtica dioica, Verbascum densiflorum* and *Veronica maritima*). We quantified plot-level pubescence as the summed cover of these five species. Data about the mean area of individual leaves were obtained from TRY – a global database of plant traits (53) TRY version 5.0; data used of (54–65). Three species were not included in the leaf area calculation, since they either had no leaves (*Cuscuta europaea* and *Equisetum pratense*) or because there were not data available in the TRY database (*Carex praecox*). All three species occurred on a maximum of two plots, and in these, with densities below 5 % cover. For an estimate of the leaf area per plot, we summed the cover-weighted leaf areas of all species per plot.

### Data analysis

All statistical analyses were done with the statistical software R (46). We ran two separate linear models to investigate which factors drove sedimentation on the vegetation and on the sediment traps. We also calculated the ratio of sedimentation on the vegetation to the sedimentation on the traps and run a separate linear model to explain it. Further, we ran six linear models to explain total amount of C, N and P in the sediment on the vegetation and in the sediment on the traps. We used the candidate variables listed in Table 1 as explanatory variables; in the ratio model we additionally included the interaction of the river kilometre and the hydrological connectivity, while in the C, N and P models, we additionally used the sediment amount as an explanatory variable. To meet model requirements regarding the normality of the error distribution, the two variables, “sedimentation on traps” (except for the ratio of sediment on vegetation to on traps) and “hydrological distance”, were natural log-transformed. We scaled all continuous variables to ensure comparability of the model estimates. To avoid multicollinearity, we removed explanatory variables with a variation inflation factor above 5.0 (vif function, car library, 66). With the remaining variables, we selected the final model with best model fit based on Akaike’s Information Criterion (stepAIC function, MASS library, 67). We tested the differences of the N:P ratios close and far from the river using paired two-sample t-tests. Therefore, the plots were separated by the mean of the hydrological distance.

## Results

### General results

The median sedimentation on the vegetation was 28.60 g m^-2^, while on the traps the median sedimentation was about double (60.55 g m^-2^, Table 2). Both, sedimentation on the vegetation and on the traps were highly variable. Sedimentation on vegetation ranged from 10.36 to 105.56 g m^-2^ and sedimentation on traps even ranged from 4.25 to 4955.50 g m^-2^, where some sediment traps that were heavily packed with sediment (Table 2). Descriptive statistics for C, N and P and for the explanatory variables are shown in Table 2.

**Table 2:** Descriptive statistics. Descriptive statistic of all continuous variables. Min=minimum, Max=maximum, Sd=Standard deviation. * the length is standardized between the images, however not calibrated to any unit.

| Variables | Unit | Min | Max | Mean | Median | Sd |
| --- | --- | --- | --- | --- | --- | --- |
| Sediment on vegetation | g m<sup>-2</sup> | 10.36 | 105.56 | 37.33 | 28.60 | 25.96 |
| Sediment on traps | g m<sup>-2</sup> | 4.25 | 4955.50 | 832.57 | 60.55 | 1440.33 |
| C in sediment on vegetation | g m<sup>-2</sup> | 0.82 | 18.79 | 4.67 | 3.76 | 3.88 |
| N in sediment on vegetation | g m<sup>-2</sup> | 0.05 | 1.00 | 0.37 | 0.36 | 0.22 |
| P in sediment on vegetation | g m<sup>-2</sup> | 0.01 | 0.28 | 0.10 | 0.09 | 0.07 |
| C in sediment on traps | g m<sup>-2</sup> | 0.56 | 178.49 | 26.09 | 3.98 | 42.68 |
| N in sediment on traps | g m<sup>-2</sup> | 0.04 | 12.88 | 1.88 | 0.30 | 3.06 |
| P in sediment on traps | g m<sup>-2</sup> | 0.02 | 3.78 | 0.87 | 0.16 | 1.09 |
| Vegetation cover | % | 7.90 | 90.20 | 50.77 | 52.61 | 21.31 |
| Biomass | g m<sup>-2</sup> | 30.12 | 499.16 | 239.51 | 219.36 | 116.20 |
| Vertical density | % | 0.08 | 0.35 | 0.20 | 0.19 | 0.05 |
| Mean height | length* | 0.09 | 0.55 | 0.25 | 0.20 | 0.10 |
| Median height | length* | 0.09 | 0.55 | 0.24 | 0.21 | 0.10 |
| Height variation | length* | 0.01 | 0.18 | 0.06 | 0.03 | 0.05 |
| Highest leaf 16 | cm | 36.00 | 124.00 | 72.08 | 73.00 | 26.00 |
| Highest inflorescence 16 | cm | 0.00 | 141.00 | 66.17 | 75.50 | 43.87 |
| Highest leaf 17 | cm | 16.00 | 72.00 | 31.25 | 23.00 | 16.37 |
| Highest inflorescence 17 | cm | 0.00 | 91.00 | 14.67 | 0.00 | 29.61 |
| Hydrological distance | m | 2.83 | 586.13 | 142.53 | 91.82 | 156.82 |
| Elevation above river | m | 0.26 | 1.71 | 1.24 | 1.31 | 0.37 |
| River kilometre | km | 3.64 | 6.98 | 5.15 | 4.99 | 1.08 |
| Shannon diversity index |  | 0.14 | 1.73 | 1.12 | 1.16 | 0.44 |
| Leaf pubescence | % | 0.00 | 37.50 | 6.90 | 2.50 | 9.27 |
| Leaf area | cm<sup>2</sup> | 234.29 | 3906.17 | 1487.25 | 1602.99 | 879.92 |

### Sedimentation on and underneath the vegetation

Sedimentation on the vegetation was influenced most strongly by the amount of vegetation biomass, but also by log hydrological distance and the height variation of the vegetation as well as the river kilometre (R^2^=0.56, Table 3). The amount of sediment on the vegetation increased with increasing biomass (*p*<0.01; Fig 2a) and decreased with increasing height variation of the vegetation (*p*=0.03; Fig 2b). In addition, sedimentation on the vegetation decreased with log hydrological distance from the river (*p*=0.01; Fig 2c), while it increased with the river kilometre (*p*=0.02; Fig 2d).

**Table 3.** Model results. Statistical model results of the sedimentation on the vegetation and on the traps.

| Sediment on vegetation |  |  |  |  |  |
| --- | --- | --- | --- | --- | --- |
|  | Estimate | Std. Error | t value | Pr(>\|t\|) | Sig |
| (Intercept) | 37.3320 | 3.5080 | 10.6420 | 0.0000 | *** |
| River kilometre | 9.5700 | 3.8320 | 2.4970 | 0.0231 | * |
| log Hydrological distance | -12.0610 | 4.4330 | -2.7210 | 0.0145 | * |
| Biomass | 14.4820 | 3.9990 | 3.6220 | 0.0021 | ** |
| Highest inflorescence 16 | -6.8990 | 5.0780 | -1.3590 | 0.1920 |  |
| Vertical density | 7.4390 | 3.8380 | 1.9380 | 0.0694 | . |
| Height variation | -9.6850 | 4.0560 | -2.3880 | 0.0288 | * |

| Sediment on trap |  |  |  |  |  |
| --- | --- | --- | --- | --- | --- |
|  | Estimate | Std. Error | t value | Pr(>\|t\|) | Sig |
| (Intercept) | 5.7200 | 0.5990 | 9.5490 | 6.83E-09 | *** |
| River kilometre | -0.7547 | 0.4264 | -1.7700 | 0.0920 | . |
| log Hydrological distance | -1.4044 | 0.3458 | -4.0610 | 0.0006 | *** |
| Precipitation | -1.4622 | 0.8481 | -1.7240 | 0.1001 |  |

**Fig 2.**
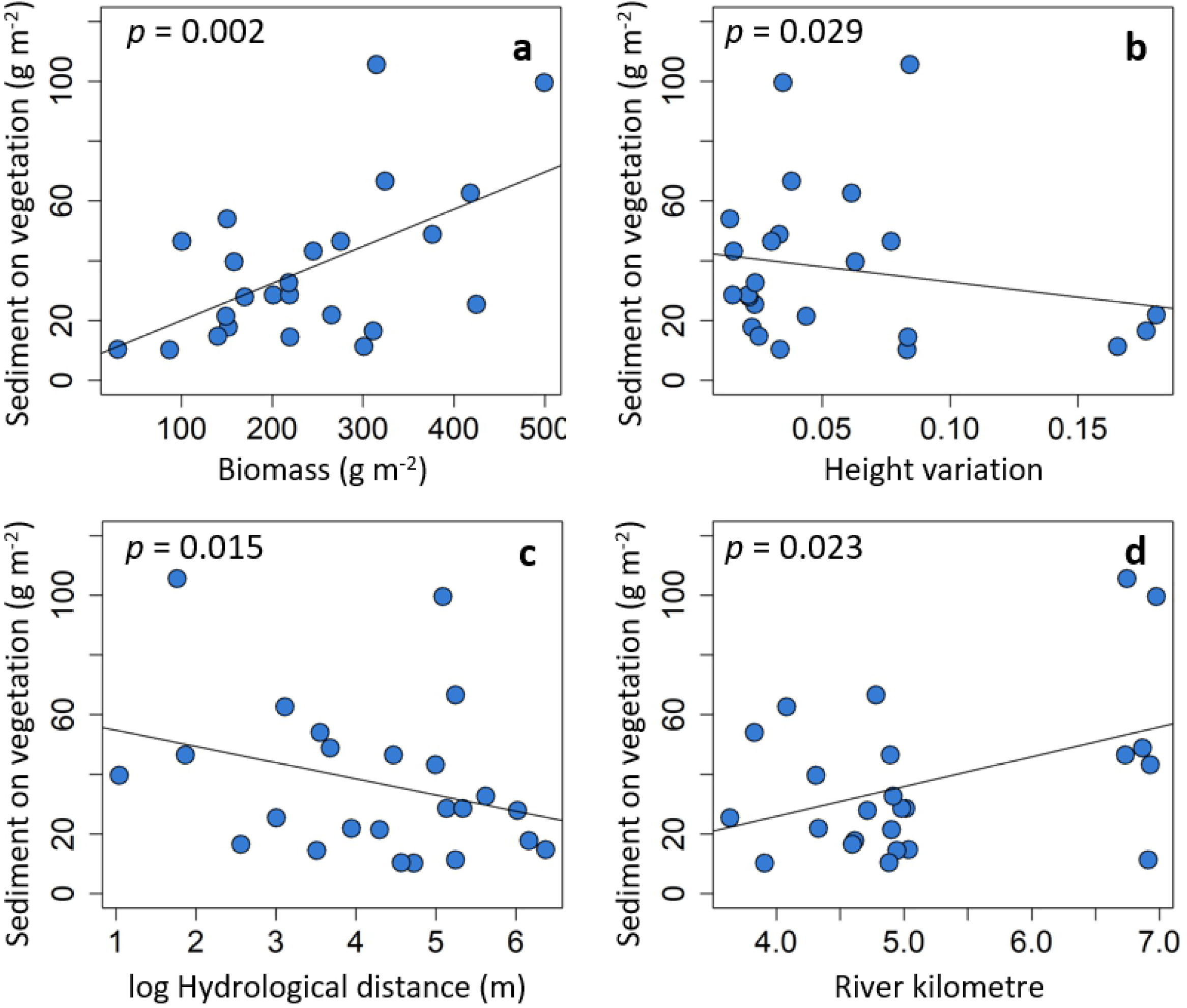
Sedimentation on the vegetation. Sedimentation on the vegetation explained by (a) biomass, (b) height variation, (c) log hydrological distance, and (d) river kilometre.

The sedimentation on the sediment traps was driven by a single topographic variable, the log hydrological distance to the river (R^2^=0.42, Table 3). Sediment traps with a short hydrological distance (close to the river) collected more sediment, and sedimentation decreased with a larger hydrological distance (*p*<0.01, Fig 3).

**Fig 3.**
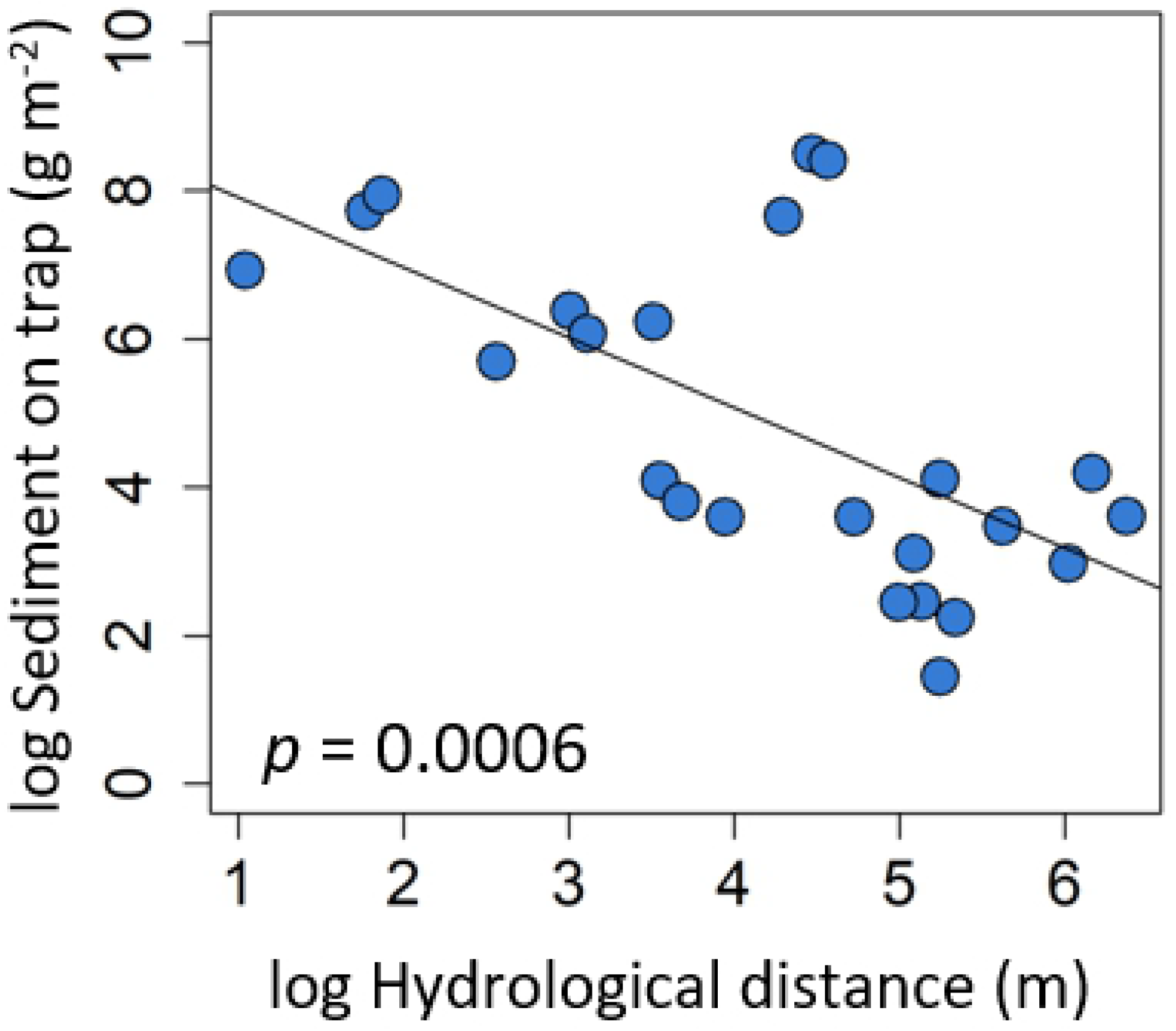
Sedimentation on traps. Sedimentation on traps explained by log hydrological distance.

Additionally, the ratio of sedimentation on the vegetation to sedimentation on the traps was driven by the hydrological distance and, the river kilometre as well as their interaction (R^2^=0.62, S1 Table). The ratio was low with short hydrological distance, meaning that relatively more sediment settled on the traps close to the river, and decrease with increasing hydrological distance (p<0.01, S4a Figure). There was also relatively more sediment on the traps at the upstream study sites, while sedimentation on the biomass relatively increased downstream the river (p<0.01, S4b Figure). The interaction of river kilometre and hydrological distance was also significant (p<0.01, S4b Figure), showing that with increasing river kilometre (i.e. more downstream), the relative increase of sedimentation on the vegetation is stronger with hydrological distance than at more upstream sites.

### Carbon, nitrogen and phosphorus in the sediment

Carbon, nitrogen and phosphorus content in the sediment strongly increased with the total amount of sediment on the vegetation (Fig 4) and log sediment on the traps (*p*<0.01 for all models, S2 Table). In addition, N on the vegetation increased with vegetation biomass (*p*=0.01) and with log hydrological distance (*p*<0.01, R^2^=0.79, Fig 4, S2 Table). Carbon and P on the vegetation additionally increased with log hydrological distance (both *p*<0.01, R^2^=0.80 and 0.92, respectively, Fig 4, S2 Table). Carbon and N content in the sediment on the traps increased with the river kilometre (both *p*=0.02, R^2^=0.64 and 0.62, respectively, S2 Table), while P content in the sediment on the traps was only explained by the amount of sediment on the trap (R^2^=0.84, S2 Table).

**Fig 4.**
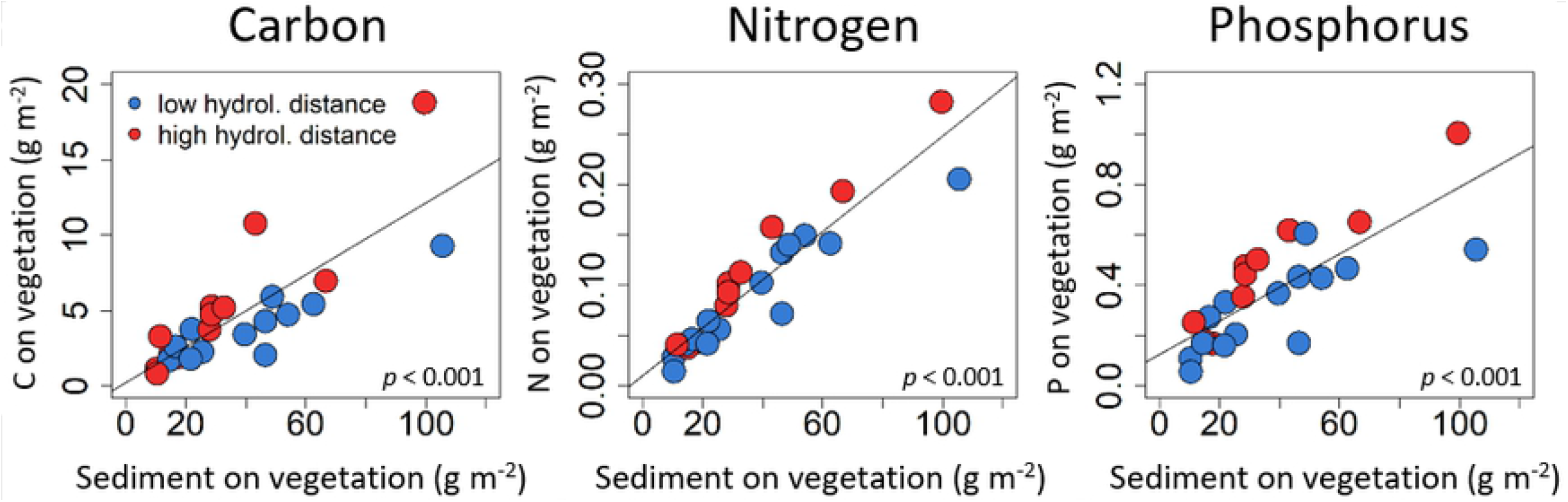
Nutrients on the vegetation. Carbon, nitrogen and phosphorus on the vegetation explained by the amount of sediment on the vegetation, and grouped for low and high hydrological distances from the river.

The N:P ratio in the sediment on the vegetation for sites closer to the river and further away from the river did not differ significantly (*p*=0.095). However, there was a trend towards a higher N:P ratio further away from the river. The same comparison (close and far away from the river) for the N:P ratio in the sediment on the traps showed a significantly higher N:P ratio for the sites further away from the river (*p*=0.001).

## Discussion

With this study, we disentangled *in situ* measurements of sedimentation on and underneath the vegetation on a floodplain and quantifying its relative importance in relation to topographic drivers. Biomass and height variation increase sedimentation on the vegetation, while vegetation characteristics did not explain sedimentation underneath the vegetation. The hydrological distance was a key variable explaining sediment and nutrient retention on and underneath the vegetation. Carbon, N and P on the vegetation increased with hydrological distance from the river in spite of the decreasing amount of sediment with increasing hydrological distance. We could not find evidence that species diversity and leaf surface structure affect the amount of sediment and nutrient retention.

### Vegetation characteristics

Regarding hypothesis (H1), we found evidence that sedimentation on the vegetation increased with increasing plant biomass and decrease with height variation. More vegetation biomass is able to provide a larger surface for sediment to settle, and thus increase sedimentation on the biomass, as it was found in the flume experiments (23,26). However, we also expected that the sedimentation on the ground underneath the vegetation would increase with increasing biomass as a consequence of a stronger reduction in flow velocity, as it was found in the flume experiment (23), but this was not supported by our findings. Three reasons might explain this: (1) it is likely that larger grain sizes (sand) accumulated underneath the vegetation, which might be less affected by the biomass above; (2) the effect of the hydrological distance on the sedimentation underneath the vegetation overrides the effects of the vegetation structure; and (3) decomposition of the plant biomass started and might already change the vegetation structure compared to the flume experiment conducted at the biomass pike. Other studies found positive or nonsignificant relations between standing biomass and trapped sediment on the ground (20,24,38). In general, we expect sediment on the vegetation to be finer grained (silt and clay), since larger grain sizes (sand or coarser) do not adhere on most of vegetation surfaces. Many important plant nutrients occur in or are associated with fine sediment (40,68). Thus, this clearly shows (1) the relevant role of standing biomass for sediment retention during the flood season, and (2) emphasizes the importance of the vegetation surface for fine sedimentation and nutrient retention.

In the flume experiment it was found that density increases sedimentation on the vegetation (23), which only showed a marginally significant increase in the present study. We did not find any statistical evidence that the vegetation height explains sedimentation, but other studies did (22,23,30). However, we found that variation of vegetation height explained sedimentation on the vegetation, even though most of the vegetation was not fully inundated. The stronger the height variation, the lower was the sediment retention on the vegetation, meaning that a more even vegetation surface collected more sediment on the vegetation. The same was found in the flume experiment (23). Others found that the intercepted biovolume calculated by the vegetation cover times the inundation depth explained a large fraction of the sedimentation on the ground (69). We could not measure the inundation depth (water level above the ground per plot), which we expected that it would increase the importance of the vegetation height and density.

### Topography

Regarding topographic parameters, we found support for hypothesis (H2) that sedimentation on the vegetation as well as underneath the vegetation decreased with increasing hydrological distance to the river. In contrast, C, N and P on the vegetation increased with the hydrological distance.

With increasing distance from the river, the flow velocity is likely to decrease and more sediment has already settled, thereby reducing the potential sedimentation on plots with longer water paths. Even though decreasing sedimentation on and underneath the vegetation was observed with hydrological distance, the three plots farthest away from the river did not had the lowest sedimentation rates; they were more than 400 m (413 - 586 m) away, while all other plots were in the range of 300 m to the river. In the same three plots the sedimentation, especially underneath the vegetation, was still reasonably high (19.65 – 66.85 mg m^-2^ [overall median 60.55 mg m^-2^]), which is in contrast with other studies that found exponential decreasing sedimentation rates on horizontal lines in the floodplain (70,71). Also other studies found decreasing amounts of sediment with increasing straight distance from the river (27,29,40,72), with increasing flow path (42) and with decreasing hydrological connectivity (15,43,44). Our result show the substantial role of shallow sites, such as abandoned meander and depression within the floodplain for sediment retention. We additionally found that the ratio of sedimentation on vegetation and on the traps increased with hydrological distance. Thus, our results emphasize the crucial role of vegetation for floodplain sedimentation.

With increasing river kilometre sediment on the vegetation and C and N underneath the vegetation increased. We expected that all three study areas receive comparable amounts of sediment with respect to quality and quantity. However, it is possible that the sites further downstream (further away from the last tributary) receive less sediment with larger grain size than the ones further upstream. We also visually observed lower flow velocities at the downstream site, at least for those plots close to the stream, which might additionally cause hither fine grained sediment, C and N retention with increasing river kilometre. For a better understanding of the key drivers, more hydraulic and hydromorphological parameters, such as discharge, inundation duration and flow velocity need to be included in the analysis (71). Still, while results could have been different for e.g. more extreme floods, our study helps to improve our general understanding of the mechanisms and processes causing sedimentation on floodplains.

### Carbon, nitrogen and phosphorus on the vegetation

Our results further support the hypothesis (H5) that nutrients (C, N and P) in the sediment increased with the amount of sediment. In addition to that, this study shows that C, N and P on the vegetation increased with greater hydrological distance. Thus, we observed relatively more nutrients on the vegetation far away from the river even though there is less total amount of sediment. Carbon and P are bound to fine grained sediment, while nitrogen is only partially associated with sediment, but it still follows similar distribution patterns (40,73). Thus, we can derive that the vegetation primarily captures finer sediment (silt, clay, and organic material), which probably also decreases in size with distance from the river, but has more nutrients bound to it. With this result, our study emphasized again the crucial role of shallow sites far inside the floodplain, such as abandoned meander and depression, for fine sediment and nutrient retention during floods.

In addition, we found an increasing N:P ratio for sites further away from the river. These changes in elemental ratios provided evidence of changes in the nutrient composition of the sediment with distance to the river main channel. A higher N:P ratio indicated a higher N availability compared to P, which suggests that N is relatively more limiting for plant growth close to the river channel, and that P is relatively more limiting for plant growth further away from the river main channel. Subsequent mineralization processes could provide additional nutrient sources for plant growth and stimulate nutrient uptake in terrestrial parts of the floodplain, as well as it might also affect community composition due to changed availability of plant nutrients (74).

### Diversity and leaf surface structure

We did not find any evidence for our hypotheses regarding species diversity (H3). The flume experiment also only showed effects of species richness on sedimentation, when species identity effects were not considered (26). Similarly, others did not find any significant differences in sediment capture capacity between monocultures and a three-species mixture in an experiment (37). Nevertheless, it is known that species diversity can correlate with vegetation structure (34), and in the flume experiment it was found that structural diversity increase sedimentation on patches (23). From grassland experiments we know that more diverse vegetation is denser and taller than low diverse vegetation (35,36).

We also did not find evidence for the importance of the leaf pubescence and leaf area in this study (H4), even though in previous studies both have been found to represent relevant traits for sedimentation (38,39). Three reasons might explain that: (1) Pubescent species were rather poorly represented within our floodplain (five species with a cover mean of 6.9 %), so that we had limited statistical power to test for its potential effects. (2) Including stem density and mean number of leaves per individual seems likely to allow a more precise estimation of the pubescence and the leaf area effect at the plot level (38). (3) Especially for leaf pubescence the seasonality of the flood could be relevant, since decomposition processes might already have diminished the leaf hairs.

### Conclusion

With our *in situ* measurements, we improve the understanding of sediment and nutrient retention in floodplains by providing insights on the vegetation structure besides the floodplain topography and simultaneously disentangling sedimentation on and underneath the vegetation. Notably, we found that more biomass increases sediment and nutrient retention on the vegetation. Sedimentation decreases with hydrological distance to the river, even though it is still reasonably high beyond distances of 400 m. Nutrients (C, N, and P) in the sediment on the vegetation, however, increase with distance to the river. Based on the results about sediment and nutrient retention, we can recommend the following management practices: First, reduced mowing for more standing vegetation biomass during the flood season, since biomass increase sediment and nutrient retention. Especially, for nutrient retention, this counts for shallow areas with high hydrological distance to the river. The mowing regime might be less important, if the focus is on maximal sediment retention, which on a mass basis happens more strongly underneath the vegetation without clear effects of the vegetation structure. Of course, trade-offs between sediment retention and other management goals, such as biodiversity conservation, should be taken into consideration when making decisions about floodplain management. Second, the strong importance of the topographical variable ‘hydrological distance’ for sediment and nutrient retention emphasizes the high value of laterally connected river-floodplain systems, including long abandoned meanders and depressions. Thus, our study suggests (1) an improvement of lateral connectivity to be able to use the potential retention hotspots far inside the floodplain, and in accordance with that (2) an adapted mowing regime on the floodplain to achieve the management regarding sediment and nutrient retention, and therefore the ecosystem function of water purification of the river.

## Acknowledgments

We thank the Federal Ministry of Education and Research (BMBF) and the Federal Agency for Nature Conservation (BfN) for funding the project (Wilde Mulde - Revitalisation of a dynamic riverine landscape in Central Germany (funding label: 01LC1322E)). We thank the TRY-initiative - a global database of plant traits and all contributors for providing the data. We also thank our student assistants (Antonia Ludwig and Georg Rieland) for supporting the fieldwork and Katie Barry for English prove reading.

## Supporting information

**S1 Table: Model results.** Statistical model results of the ratio sediment on the vegetation to sediment on the traps.

**S2 Table: Model results.** Statistical model results of carbon, nitrogen and phosphorus on the vegetation and on the traps.

**S1 Figure: Map of the study site.** Map of the three floodplains along the Mulde River with trap locations.

**S2 Figure: Structural photo.** a) Original photo with blue background wall and blue flooring material in front. b) Automatically analyzed images for vertical density and height distribution (done with R) with sketch of variables calculated from the image.

**S3 Figure: Sediment traps.** Picture of a sediment trap in the field.

**S4 Figure: Sedimentation ratio.** Ratio of sediment on vegetation to sediment on traps.

